# Seeing what you hear: compression of rat visual perceptual space by task-irrelevant sounds

**DOI:** 10.1101/2025.02.11.637608

**Authors:** Mattia Zanzi, Francesco G. Rinaldi, Silene Fornasaro, Eugenio Piasini, Davide Zoccolan

## Abstract

The brain combines information from multiple sensory modalities to build a consistent view of the world. The principles by which multimodal stimuli are integrated in cortical hierarchies are well studied, but it is less clear whether and how unimodal inputs systematically shape the processing of signals carried by a different modality. Here we use a visual classification task in rats to investigate how task-irrelevant sounds modify the processing of visual stimuli. Our data shows that the intensity of a sound, but not its temporal modulation frequency, enacts a powerful, effective compression of the visual perceptual space. This result underscores the importance of cross-modal influences in perceptual pathways and suggests an important role for inhibition as the mediator of auditory-visual interactions at the neural representation level.

---

In the cerebral cortex, integration of information from different sensory modalities takes place at multiple stages along the sensory pathways. The process that is best understood at the functional level is the way unisensory cues can be combined by high-order association cortices, often in a statistically optimal way, to increase the accuracy of perceptual judgements (*1–8*). By comparison, the functional impact on perception of the hetero-modal inputs that reach primary sensory cortices in both primates (*9–12*) and rodents (*13–18*) is less well established.

For instance, several authors have found direct cortico-cortical connections from primary auditory (A1) to visual (V1) cortex in rodents (*13–16*), but studies disagree on the impact of these projections on local cortical dynamics and encoding of visual stimuli. Sound was originally reported to hyperpolarize supragranular pyramidal cells in mouse V1, by recruiting a translaminar GABAergic network activated by A1 projections to infragranular V1 neurons (*14*). Later studies, however, found that most A1 projections terminate in superficial layers of V1 and engage local inhibitory and disinhibitory circuits underlying a much richer variety of auditory influences on V1: enhancement and suppression of visually evoked responses have both been documented, as well as sharpening of orientation tuning and improvement in the coding of visual information (*13, 15, 17, 18*). This heterogeneous assortment of sound-mediated modulations in V1 seems to depend on factors such as the contrast of the visual stimuli, the luminosity of the environment, the spectral (e.g., pure tones vs. noise bursts) and envelope (e.g., loud vs. quiet onset) properties of the sounds and the temporal congruency between visual and auditory stimuli. Finally, it remains highly debated the extent to which, in awake mice, sound-driven responses in V1 reflect auditory inputs or, rather, behavioral modulation via sound-evoked orofacial movements (*16, 17, 19*).

Amid such contrasting findings at the cortical circuitry level, the functional impact of auditory signals on rodent visual perception is poorly explored. Does sound improve acuity in visual discrimination tasks? Can any signature of sound-induced suppression (or enhancement) of visual representations be found at the behavioral level? Is the temporal consistency between sounds and visual stimuli important in this regard?

To address these questions, we trained 10 rats in a visual temporal frequency (TF) classification task, where the visual (V) stimuli to categorize were paired with simultaneous but task-irrelevant auditory (A) stimuli (Fig. 1A; see also fig. S1). The visual stimuli were circular sinusoidal gratings moving outward with 9 possible, randomly selected TFs (Fig. 1B; x axis). Rats categorized these stimuli as “Low TF” or “High TF”, relative to an ambiguous (randomly rewarded) boundary value (2.12 Hz) in the middle of the tested TF range (vertical dashed lines in Fig. 1C and D). The auditory stimuli consisted of either fixed amplitude or amplitude modulated (AM) white noise bursts (fig. S2A). The sounds were uninformative about the correct classification of the visual stimuli, to avoid high-level multisensory integration effects and focus instead on the direct impact of auditory signals on the representation and perception of visual stimuli.

**Figure 1:**
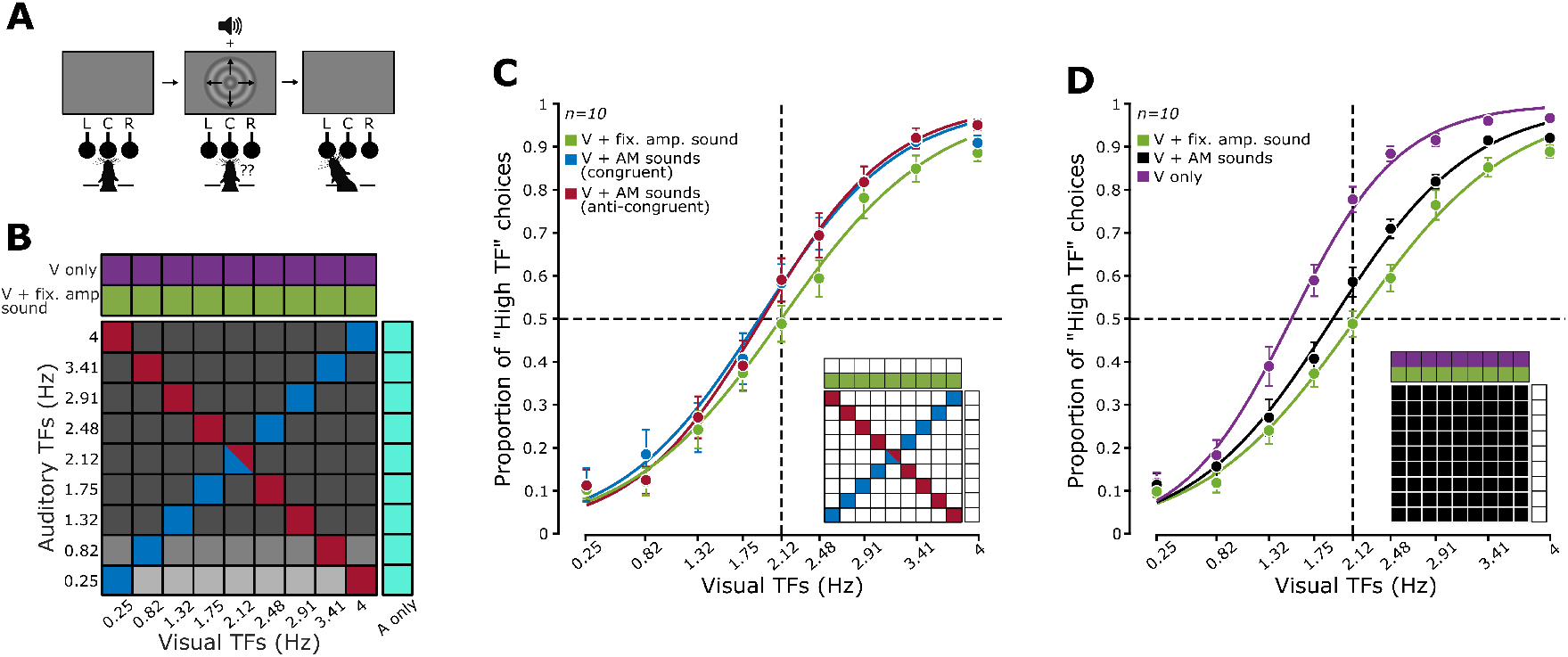
Impact of sound on rat classification of visual temporal frequencies. (**A**) Schematic representation of a behavioral trial. Rats were trained in sound-isolated operant boxes equipped with a display and a speaker for presentation of visual and auditory stimuli (see also fig. S1). As in previous studies (*21, 22*), the rats learned to protrude their head through an opening in the box, so as to face the display and the speaker and trigger stimulus presentation by touching the central nose-poke port in a three-port array. The visual stimuli were circular, outward-moving (see arrows) sinusoidal gratings that rats learned to classify according to their TF. Gratings with TF ¡ 2.12 HZ had to be classified as “Low TF”, while those with TF > 2.12 Hz as “High TF”. Each class was associated with one of the two lateral nose-poke ports, and correct responses were rewarded with a drop of stevia-water solution. Auditory stimuli were task irrelevant. (**B**) Stimulus conditions presented to the rats during the test phase of the experiment. These included: 1) visual gratings paired with fixed amplitude sounds (green cells); 2) visual gratings paired with AM sounds (all possible combinations of TFs in the two modalities; cells in the squared matrix; blue, gray and red cells refer, respectively, to congruent, incongruent and anti-congruent audiovisual conditions; the shades of gray, from light to dark, indicate AM sounds with progressively larger intensity; see main text for details); 3) unimodal purely visual gratings (purple cells); and 4) unimodal purely auditory AM sounds (cyan cells). (**C**) Group average proportion of “high TF” choices (*n*=10 rats) as a function of the TF of the visual gratings, when the latter were paired with: 1) fixed amplitude sounds (green); 2) congruent AM sounds (blue); and 3) anti- congruent AM sounds (red). The error bars denote s.e.m. over the group of 10 rats. The curves are the result of logistic regressions to the data points. The inset illustrates which of the stimulus conditions shown in **B** contributed to the data points/curves. Table 1 reports the results of a 2-way ANOVA having as factors the experimental condition (“V + fixed amplitude sound”, “V + AM congruent” and “V + AM anti-congruent”) and the grating TF, as well as the results of a post-hoc Tukey test between each pair of experimental conditions. (**D**) Same as in **C**, but for stimulus conditions where the gratings were paired with: 1) fixed amplitude sounds (green; same data as in **C**); 2) AM sounds (black); and 3) no sounds (purple). Table 2 reports the results of a 2-way ANOVA having as factors the experimental condition and the grating TF, and the results of a post-hoc Tukey test between each pair of experimental conditions.

During the training phase, the gratings were paired with the constant amplitude noise burst. Rats were gradually introduced to the TFs they had to classify, starting from the most discriminable ones (0.25 vs. 4 Hz), until they reached an average performance higher than 70% correct over the non-ambiguous TF values (fig. S3). The contrast of the gratings was initially set at 100%, but it was progressively reduced to 25%, given that the impact of sound on V1 responses has been found to be stronger when visual stimuli are shown at low/intermediate contrast (*13, 17*). Once the animals had become proficient in this task (fig. S3), they were moved to the testing phase, where they faced a rich variety of audiovisual stimuli (illustrated in Fig. 1B). The gratings were paired not only with the fixed amplitude burst used during training (Fig. 1B, green cells), but also with AM white noise stimuli. These sounds were obtained by modulating the amplitude of a white noise burst with sinusoids having 9 possible TFs (the same of the visual gratings). The rats were presented with all possible pairwise combinations of visual and auditory TFs (cells in the squared matrix of Fig. 1B). This yielded trials where the auditory and visual stimuli had either the same TF (blue cells) or different TFs (gray and red cells). The latter included a special pool of anti-congruent conditions (red cells), where progressively larger visual TFs were paired with progressively smaller auditory TFs. Additionally, we also tested unimodal, purely visual trials (purple cells) and unimodal, purely auditory trials with AM sounds (cyan cells). Since the rats were never trained to classify the TFs of the auditory stimuli, with the latter trials we simply measured the spontaneous responses of the animals to the AM sounds, without providing any feedback about their choices (*20*). The rats did not spontaneously classify the AM bursts according to their TF in the auditory only conditions, thus confirming the task-irrelevance of the sounds (fig. S4).

Overall, this design allowed probing the impact of task-irrelevant auditory stimuli on visual perception along two distinct dimensions: 1) whether sound alters the sensitivity of visual discrimination or, rather, introduces a bias in visual classification; and 2) whether the impact of sound depends on the temporal consistency of audio-visual stimuli or, rather, on the intensity of the auditory stimuli.

When the gratings were paired with the fixed amplitude noise, the group average psychometric curve reporting the proportion of “high TF” choices as a function of the grating frequency was sharp and symmetrical, with the point of subjective equality sitting squarely on the ambiguous TF (Fig. 1C, green). Interestingly, pairing the gratings with either the congruent (blue) or anti-congruent (red) AM sounds produced an identical increase in the proportion of “high TF” choices, which was particularly prominent for the highest TFs. This observation was confirmed by a 2-way ANOVA having as factors the audiovisual experimental condition and the grating TF (*20*). Both factors significantly modulated rat choices and so did their interaction (Table 1), thus confirming that the increase of “high TF” choices yielded by the modulated sounds was not homogeneous along the TF axis. A post-hoc Tukey test confirmed that the curves observed for the two conditions featuring AM sounds were statistically indistinguishable. This indicated that the congruency between the TF of the AM sounds and the TF of the gratings did not affect the impact of the auditory stimuli on the visual discrimination task.

**Table 1:** Two-way ANOVA table (a) and Tukey post-hoc tests (b) for the comparison among rat psychometric curves shown in Fig. 1C.

**(a) Two-way ANOVA with repeated measures**
| **Effect** | **df** | **SS** | **MS** | **F** | **p** |
| --- | --- | --- | --- | --- | --- |
| Visual TF | 8 | 33.483 | 4.185 | 289.291 | $0.001 \cdot 10^{-48}$ |
| Experimental condition | 2 | 0.212 | 0.106 | 4.897 | 0.02 |
| Visual TFs:Experimental condition | 16 | 0.191 | 0.012 | 2.533 | 0.0018 |

**(b) Tukey post-hoc tests**
| **Condition 1** | **Condition 2** | **Difference** | **StdErr** | **p** | **Lower** | **Upper** |
| --- | --- | --- | --- | --- | --- | --- |
| V + AM sounds<br>(congruent) | V + AM sounds<br>(anti-congruent) | -0.0069 | 0.0118 | 0.8297 | -0.0397 | 0.0259 |
| V + AM sounds<br>(anti-congruent) | V + fixed-amplitude<br>sound | 0.0625 | 0.0254 | 0.0839 | -0.0084 | 0.1336 |
| V + fixed-amplitude<br>sound | V + AM sounds<br>(congruent) | -0.0556 | 0.0256 | 0.1298 | -0.1271 | 0.0159 |

This observation led us to group together all the trials where the visual stimuli were paired with an AM sound, regardless of the congruency of their TFs. The resulting psychometric curve (Fig. 1D, black) featured an asymmetrical vertical shift, with respect to the reference curve measured with the fixed amplitude sounds (green dots/line), that was equivalent to the one previously observed with the congruent and anti-congruent AM sounds (compare to the blue and red curves in Fig. 1C). Strikingly, a similar but even more prominent shift of the psychometric curve was observed for the unimodal visual conditions, when the gratings were presented without any concomitant sound (purple dots/line). Again, a two-way ANOVA showed a significant main effect of experimental condition, TF of the gratings and their interaction, and a post-hoc Tukey test revealed a significant difference between the curve obtained with the unimodal visual stimuli and the curves measured with the paired sounds, either AM or fixed amplitude (Table 2).

**Table 2:** Two-way ANOVA table (a) and Tukey post-hoc tests (b) for the comparison among rat psychometric curves shown in Fig. 1D.

**(a) Two-way ANOVA with repeated measures**
| **Effect** | **df** | **SS** | **MS** | **F** | **p** |
| --- | --- | --- | --- | --- | --- |
| Visual TF | 8 | 33.809 | 4.226 | 314.013 | $0.002 \cdot 10^{-53}$ |
| Experimental condition | 2 | 1.766 | 0.883 | 32.593 | $0.001 \cdot 10^{-5}$ |
| Visual TFs:Experimental condition | 16 | 0.461 | 0.29 | 8.763 | $0.001 \cdot 10^{-11}$ |

**(b) Tukey post-hoc tests**
| **Condition 1** | **Condition 2** | **Difference** | **StdErr** | **p** | **Lower** | **Upper** |
| --- | --- | --- | --- | --- | --- | --- |
| V + fixed-amplitude | V + AM sounds | -0.0614 | 0.0244 | 0.0769 | -0.130 | 0.0067 |
| V + AM sounds | V only | -0.1324 | 0.1776 | 0.0001 | -0.182 | -0.0829 |
| V only | V + fixed-amplitude | 0.1938 | 0.0299 | 0.0003 | 0.1102 | 0.2773 |

These results showed that rat perception of visual temporal frequencies was systematically shifted towards reporting the “high-TF” category if the noise bursts became amplitude modulated and, even more so, when the sounds disappeared altogether. Wondering about the possible cause of these progressively larger shifts, we realized that one key attribute of the sound did decrease from fixed amplitude to AM bursts and then further to the purely visual conditions: its intensity. In fact, because of the sinusoidal modulation, there were moments during the AM bursts where the instantaneous sound intensity dropped to zero. As a result, the average sound intensity (as computed over the inferred reaction time of the animals) was lower for the AM bursts than for the fixed amplitude burst (fig. S2), besides being minimal for the purely visual stimuli (corresponding, in this case, to the environmental background noise). In a previous study (*14*), noise bursts were reported to hyperpolarize V1 neurons, with the magnitude of this inhibition increasing monotonically with their intensity. Given this observation, we hypothesized that the perceptual effect of sound is to compress visual cortical representations in a way that is proportional to its intensity.

To test this hypothesis, we developed a Bayesian ideal observer model of rat perceptual choices, resting on three assumptions: 1) that V1 neurons respond to progressively larger TFs with approximately linearly increasing firing rates, at least within the TF range tested in our study (cartoon in Fig. 2A; cyan vs. red dots); 2) that sound dampens neuronal responses, compressing them towards the origin of the V1 representational space, with the compression being stronger for higher intensity sounds (compare the green to the purple distributions in Fig. 2A); and 3) that downstream decision neurons learn an optimal discrimination boundary in the representational space under the sensory conditions they were repeatedly exposed to during training (visual gratings paired with the fixed amplitude sound; green distributions in Fig. 2A) and keep it fixed when new conditions (e.g., purely visual gratings; purple distributions) are interleaved with those used for training.

**Figure 2:**
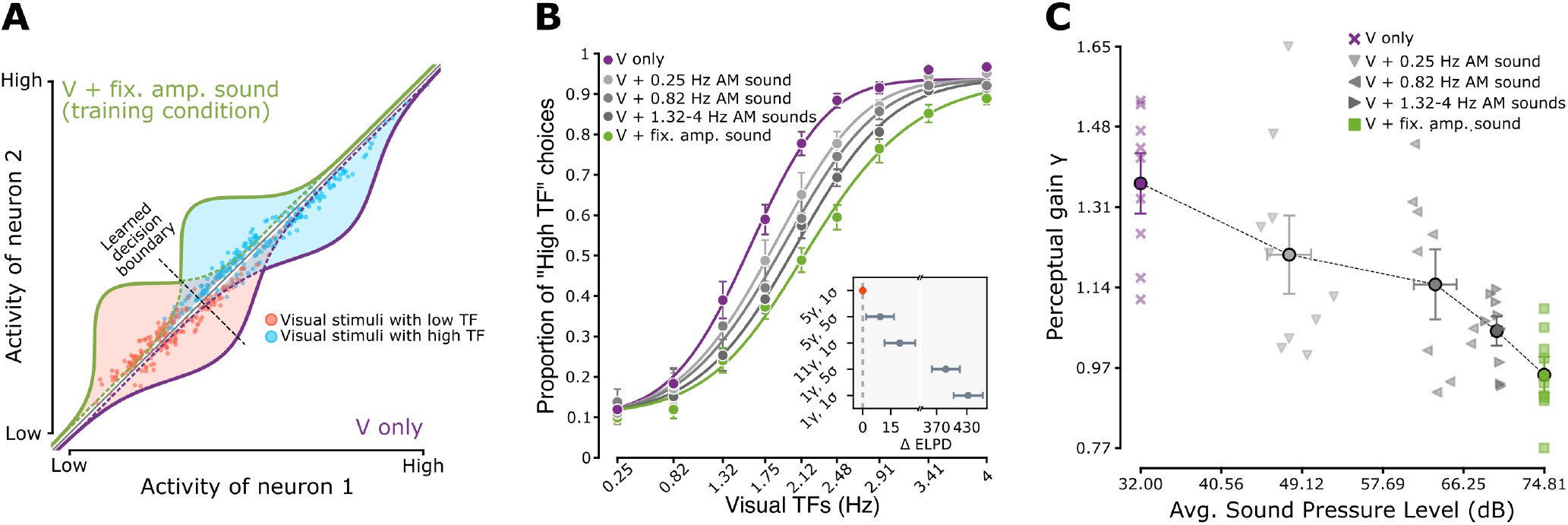
An ideal observer model where neuronal representations encode linearly visual temporal frequencies and are suppressed by sound intensity accurately accounts for rat perceptual choices. (**A**) Schematic of the assumptions at the base of the ideal observer model. The drawing illustrates a hypothetical neuronal representational space made of two units. Each neuron responds to visual gratings with an average firing rate that grows linearly as a function of the grating TF. Due to neuronal noise, the responses to repeated presentations of the same grating are distributed according to a Gaussian function centered on the average firing rate elicited by the grating. In the example, two TFs yields two distributions of population vectors (red and cyan dots). When the gratings are paired with a sound with strong intensity (e.g., the fixed amplitude noise burst), all responses are inhibited and, as a result, the distributions are compressed toward the origin of the representational space (green curves). Since rats are trained to discriminate the gratings under these conditions, they learn a decision boundary (dashed line) that optimally separates such compressed distributions. When the animals are presented with unimodal, purely visual gratings, the sound-induced inhibition is released and the response distributions shift towards higher firing rate values (purple curves). The animals, however, still rely on the previously learned decision boundary to discriminate these stimulus conditions. As a result, the overall proportion of “high TF” choices increases (i.e., more red dots fall on the “false alarm” side of the decision boundary). (**B**) Group average proportion of “high TF” choices (*n*=10 rats) as a function of the TF of the visual gratings, when the latter were paired with: 1) fixed amplitude sounds (green); 2) AM sounds with increasingly larger TFs (progressively darker shades of gray, matching those in the stimulus matrix shown in Fig. 1B); and 3) no sounds (purple). The error bars denote s.e.m. over the group of 10 rats. The curves are group-level predictions of the proportions of “high TF” choices according to the Bayesian ideal observer model defined in Equation 1, with a single value of the sensitivity parameter σ and 5 different values of the scaling factor *γ*, one for each level of sound power (more precisely, each curve is the mean of the distribution of psychometric curves induced by the posterior distribution of the σ and *γ* parameters obtained for a given sound power level). The inset shows the difference between the expected log predictive density (ELPD) of the reference model with 1 σ and 5 *γ* values (i.e., the one yielding the psychometric curves) and variants of the model having a given number of free σ and *γ* parameters (as indicated on the ordinate axis). In all comparisons, the reference model was the one with lowest ELPD, i.e., with the higher predictive power. (**C**) Relationship between the magnitude of the scaling factor *γ* in the ideal observer model and the intensity of the sounds that were paired with the visual gratings. Triangle, square and cross symbols refer to individual rats, while circles are group averages. Sound conditions are labeled according to the same color code as in **B** and Fig. 1B. The error bars denote s.e.m. over the group of 10 rats.

In our model, on each trial, the combination of a visual and an auditory stimulus generates an internal representation sampled from a Normal distribution with fixed standard deviation σ, centered on some average representation of the TF value of the visual stimulus *s*. Following from the assumptions above, the internal representation is then scaled by a factor *γ* (*20*), and the decision rule is the one that would be Bayes-optimal in the training condition. We can thus derive the probability for a rat of classifying a given stimulus *s* as belonging to the *H* (i.e., “High TFs”) category as

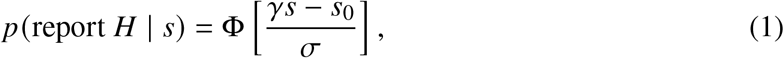

where Φ is the standard Normal cumulative function and *s*_0_ is the boundary 2.12 Hz visual stimulus that separates the “Low TFs” class from the “High TFs” class (*20*). In the model, the σ parameter controls the slope of the psychometric function and measures the internal noise of the representation (i.e., rat sensitivity over the visual TF axis), while the *γ* parameter controls the gain of the internal perceptual representation.

This model allowed testing alternative hypotheses about the impact of sounds on visual representations, depending on the number of distinct values that the σ and *γ* parameters were free to take. At one extreme, allowing a different value of σ for each sound condition would represent the hypothesis that sound affects the sensitivity of the visual representation (i.e., the sharpness of the psychometric curves). At the other extreme, keeping a constant σ but allowing a different value of *γ* for each level of sound intensity would represent the hypothesis that sound compresses the visual representation according to its intensity, but leaves visual perceptual sensitivity unaltered.

Using the model to test for the impact of sound intensity on visual perception required estimating the sound intensity experienced by the rats in the time interval between stimulus onset and each animal’s estimated reaction time (*20*) (see fig. S2B). Analyzing the resulting, trial-averaged intensity levels revealed a finer-grain grouping of stimulus conditions, compared to the one shown in Fig. 1D. In addition to the conditions with maximal and minimal intensity (i.e., 74.8 dB for the fixed amplitude sounds and 32 dB for the purely visual stimuli, respectively), we found that the AM conditions could be grouped into three main clusters with increasingly larger intensity, based on the TF at which the noise bursts were modulated: TF = 0.25 Hz (48.5 ± 2.4 dB; mean ± s.e.m. across rats); TF = 0.82 Hz (64.2 ± 2.3 dB); and 1.32 Hz ≤ TF ≤ 4 Hz (70.7 ± 0.5 dB; these conditions are shown with progressively darker shades of gray in the stimulus matrix of Fig. 1A and in Fig. 2B-C; see fig. S2C for details).

We inferred the parameters of the ideal observer using a hierarchical Bayesian approach (*20*) (see fig. S5), which yielded a posterior probability distribution over the values σ and *γ* that characterized each specific rat (referred to as rat-level parameters), as well as the values that characterized all rats taken together (referred to as group-level parameters). The group-level estimates obtained for a model with a single σ and 5 different *γ* per rat (i.e., one *γ* for each of the 5 levels of sound power in our stimulus set) are summarized by the psychometric curves shown in Fig. 2B.

The curves fitted tightly the measured classification accuracies and captured well the way the proportion of “high TF” choices progressively increased, while sound intensity dropped from maximal (green curve; fixed amplitude sound) to minimal (purple curve; no sound), passing through the three intermediate levels in which the AM sounds were clustered (gray curves). This means that an ideal observer with constant sensitivity (i.e., a single σ) could describe well the impact of sound on rat visual perceptual choices, provided that the internal representation of visual TFs was appropriately scaled by a gain factor (γ). Importantly, the magnitude of this gain decreased monotonically across the 5 levels of increasing sound intensity in which our stimulus conditions could be grouped (Fig. 2C).

To verify that this was the best description of the data, we compared the predictive power of this “reference” model to that of other variants, with a different number of values that σ and *γ* were allowed to take. The predictive power of each model was measured using the expected log predictive density (ELPD), that is, the average, expected log-likelihood of the model on new data. This metric is estimated by leave-one-out cross-validation using a standard approach and automatically penalizes overly complex models that tend to overfit (*20*). In our comparison, we computed the difference in ELPD between the reference model with 1 σ and 5 *γ* and each other model. We refer to this metric as ΔELPD.

The reference model with a fixed σ and 5 *γ* (one for each of the 5 levels of sound intensity) was the best (lowest ELPD) of a broad range of models we tested (Fig. 2B, inset). In particular, allowing a separate *γ* for each of the 9 AM conditions, thus obtaining a total of 11 possible *γ* values (while still keeping σ fixed to a single value), yielded a worse model (an indication of overfitting). This confirmed that noise bursts modulated within the [1.32 Hz, 4 Hz] range were all equivalent in terms of their impact on visual perceptual choices, as expected given their very similar intensity (fig. S2C). A model with 5 σ and 1 *γ* also performed much worse than the reference model, thus confirming that the different sound stimuli did not impact rat sensitivity to the visual TFs, but, rather, the dynamic range of the internal representation of the TFs. This conclusion was further reinforced by noticing that not even a model with 5 σ and 5 *γ* outperformed the reference model with only 1 σ and 5 *γ*.

In summary, our Bayesian ideal observer analysis supports the hypothesis that the effect of sound on the representation of temporally modulated visual gratings is to compress the visual perceptual space rather than increase or decrease perceptual uncertainty. Mechanistically, this finding suggests that sound acts as a powerful modulator of the responses of visual cortical neurons (see Fig. 2A) but leaves their tuning for temporal frequency unaltered.

This conclusion implies that, among the variety of auditory influences that have been reported in V1 (*13–18*), inhibition may play a dominant role at the functional level (*14*). At the same time, our results appear inconsistent with the sound-induced enhancement of cortical responses and the sharpening of orientation and direction tuning reported by some authors (*13, 17*). One possibility to reconcile these findings is that different kinds of perceptual tasks may be differently affected by auditory signals. In tasks requiring the discrimination of visual features that are related to the energy of the stimulus (such as its TF) and that are likely encoded by locally linear tuning curves [as postulated in our model; see Fig. 2A and (*20*)], sound-mediated engagement of inhibitory circuits may lead to a dominant, strong suppression of visually-evoked responses. In tasks where spatiotemporal features (e.g., orientation and direction) are encoded by neuronal populations with bell-shaped, unimodal tuning curves (e.g., over the orientation axis), the interplay between local inhibitory and disinhibitory circuits activated by sound could perhaps lead to a decrease of perceptual uncertainty and an improvement of decoding accuracy.

In conclusion, our study establishes that task-irrelevant auditory signals have a strong influence on the way task-relevant visual information is processed in the rodent brain. Future work on behavior and visual cortical codes will be necessary to understand the nature of this modulation and its interplay with task requirements.

## Supporting information

Zanzi et al. Supplementary Materials

## Acknowledgments

We thank Mathew Diamond, Giuliano Iurilli, Maximiliano Jose Nigro, Matteo Marsili and Carlo Fantoni for valuable discussions and insightful feedback on the manuscript.

## Author contributions

Conceptualization: M.Z., F.G.R., E.P., and D.Z. Data curation: M.Z., S.F. and F.G.R. Formal analyses: M.Z., F.G.R., E.P., and D.Z. Funding acquisition: E.P. and D.Z. Investigation: M.Z. and S.F. Methodology: M.Z., F.G.R., E.P., and D.Z. Project administration: E.P. and D.Z. Software: M.Z. and F.G.R. Supervision: E.P. and D.Z. Visualization: M.Z. and F.G.R. Writing – original draft: M.Z., F.G.R., E.P., and D.Z. Writing – review & editing: M.Z., F.G.R., S.F, E.P., and D.Z.

## Data and materials availability

In case the manuscript is accepted for publication, all source data and code needed to evaluate the conclusions of the study will be made freely available in a publicly accessible repository on Zenodo (*23*). In the meanwhile, the editor and the reviewers will be able to access this material at the following link: https://zenodo.org/records/14883380?token=eyJhbGciOiJIUzUxMiJ9.eyJpZCI6IjI5NDZkMDQyLWI4YjgtNDU4Yi05ZjRiLTc4MjdiNGYwOWMMyIsImRhdGEiOnt9LCJyYW5kb20iOiIyZGY4ZTkzNGM5ZGMzMmZlOTliMDI2NDZlZGMwNWUyYiJ9.WbBo3xG-KbMhvPMrdv-CtmjIXKmXzDyAbJR12u9uKsi0TQZbLFssULKYV4_gqom07f3nwRk7uaCIZaaOOZIO9Q

## Supplementary materials

Materials and Methods

Figs. S1 to S5

References (*25-33*)

