## Supplementary material for "Seeing what you hear: compression of rat visual perceptual space by task-irrelevant sounds": Zanzi et al. Supplementary Materials

**This PDF file includes:**

Materials and Methods

Figures S1 to S5

### Materials and Methods

#### Subjects

All animal procedures followed international and institutional standards for animal care and use in research and were approved by the Italian Ministry of Health (project approval 940/2015-PR on September 4, 2015; project approval 1183/2020-PR on December 4, 2020).

We trained 12 adult Long Evans rats (Charles River Laboratories) in a visual temporal frequency (TF) classification task. The animals, seven weeks old on arrival, weighed approximately 240 g each. They were housed three per cage in a climate-controlled chamber (23 ± 1° C, 40-50% humidity). Training began at 10 weeks, following two weeks of handling to accustom them to human interaction. Rats always had free access to food (20 g/day) but their access to water was restricted in the days of the behavioral training/test (5 days a week). Water was provided as a reward for correct responses in the discrimination task and for about 30 minutes after each session, ensuring at least the recommended 50 $\mathrm{ml\,kg^{-1}}$ daily intake. During the course of the study, their body weight increased to about 500 g. The duration of the training phase (see below) varied depending on the animal. Five rats completed it in  3 months and proceeded to the test phase (see below), while for the remaining rats it took  6 months. The test phase lasted approximately 12 months for all the rats.

#### Behavioral apparatus

Rats were trained in a rig made of two identical racks, each with three vertically stacked slots, allowing six animals to be trained simultaneously. Each slot contained a custom-built black operant chamber (20×20×30 cm) with a 4 cm-diameter viewing hole for the rats to extend their heads and face the stimulus display (Fig. S1). The visual stimuli were presented on a 21.5" LCD monitor (ASUS VE228) placed at 30 cm from the viewing hole, while the auditory stimuli were presented from a full-range speaker (Visaton BF37, 4 Ohm, 1.5" diameter, frequency response 100–20000 Hz) positioned above the monitor, tilted 45° towards the viewing hole, and located 35 cm from it (Fig. S1). Each speaker received a pre-amplified signal (Q-BAIHE TDA7377PRO). A 3D-printed block with three equidistant response ports was positioned at 3 cm from the viewing hole. Each port had a stainless-steel feeding needle and a LED-photodiode pair that worked as a licking sensor to detect interactions of the rats with the port. Signals from the photodiodes were digitized with a microcontroller (Phidgets 1203 input/output device) and sent to a computer running the open-source MWorks software (`https://mworks.github.io/`). This system automatically presented the visual and auditory stimuli when the animal's nose was detected in the central port, while the lateral ports were used to record the responses of the animals to the visual stimuli and decide whether to deliver the reward, along with reinforcement sounds (Fig. 1A). Correct responses were rewarded with a 4% stevia-water solution from the lateral feeding needles connected to computer-controlled syringe pumps (New Era Pump Systems; NE-500). Each animal was individually monitored with a camera (AXIS M1033-W Network Camera) positioned 14 cm above the chamber viewing hole. The operant chambers were acoustically insulated from each other and the environment with sound-proof foam panels.

**Visual and auditory stimuli**

Rats were presented with black-and-white outward-moving sinusoidal gratings (30 fps) with a spatial frequency of 0.05 cycles $\mathrm{deg}^{-1}$ and temporal frequencies of 0.25, 0.82, 1.32, 1.75, 2.12, 2.48, 2.91, 3.41, and 4 Hz (the temporal frequency of the gratings can be defined as the number of grating cycles moving through a point of the screen in 1 second). These frequencies were symmetrically distributed around 2.12 Hz and logarithmically spaced from this frequency (Fig. 1B, x axis). The gratings, displayed on a middle-grey background, were enveloped with a Gaussian circular aperture with diameter corresponding to the monitor height (see examples in Fig. 1A and Fig. S1).

Two classes of auditory stimuli were used in our study: fixed amplitude sounds (Fig. S2A, green curve) and amplitude modulated (AM) sounds (Fig. S2A, gray curves). In both cases, the stimuli were generated in Matlab (`https://www.mathworks.com/`) from a full-range white noise sampled at 44.1 kHz. In the case of the AM sounds, the white noise time series was used as a carrier and multiplied with positive sinusoid envelopes (the modulators), having the same temporal frequencies of the visual stimuli (Fig. 1B, y axis).

More in detail, sound stimuli were generated as digital control signals, sampled at 44.1 KHz in arbitrary units, which were then converted in WAV format (16 bits per sample). We passed the WAV files through our custom-built amplifier and loudspeaker (see "Behavioral Apparatus" and Fig. S1).

The control signal was generated starting from a white noise sequence. We used two distinct white noise sequences, one for the fixed amplitude stimulus and one for the AM stimuli. In the fixed amplitude condition, the white noise sequence (denoted as $a_{\text{fix}}(t)$) was composed of independent and identically distributed random variables sampled from a normal distribution with a sampling rate matching that of the control signal ($\nu_{\text{WN}} = 44.1$ kHz). The distribution had parameters $\mu = 0$ and $\sigma = 1$. Before conversion to WAV format the sequence was normalized to have values in the interval $[-1, 1]$, hence giving the distribution an effective standard deviation $\sigma_{eff} = 0.283$. The same sequence was used in all the "V + fixed amplitude sound" trials.

The white noise carrier of the amplitude modulated sounds, denoted as $a_{AM}(t)$, was composed of independent and identically distributed random variables sampled from a uniform distribution in an interval $[-1, 1]$, again with a sampling rate of $\nu_{\text{WN}} = 44.1$ kHz. The same sequence was used in all the trials of the modulated conditions. The sequence $a_{AM}(t)$ was multiplied by a modulation function $m(t) = \frac{1-\cos(\omega_n t)}{2}$, where $\omega_n = 2\pi\nu_n$ is the angular frequency of modulation and $\nu_n$ is the modulation frequency of condition $n$ (Fig. S2A, gray curves). The modulation was non-negative and bounded in $[0, 1]$, ensuring that the modulated signal remained within $[-1, 1]$.

**Behavioral Task**

The rats learned to initiate a behavioral trial by nose-poking the central port, and to approach the lateral ports to classify the visual stimuli based on their temporal frequency (Fig. 1A). The "low temporal frequencies" class (0.25-1.75 Hz) and the "high temporal frequency" class (2.48-4 Hz) had to be reported by half of the rats by poking respectively the left and right port, while the other half was trained with the opposite association. To initiate a trial, rats had to keep their nose in the central port for 300 ms, followed by a 100 ms tone (500 Hz pure tone) signaling the stimulus presentation, occurring 200 ms later. After stimulus onset, rats were given 2 seconds to make a response; responses within 300 ms were aborted to prevent impulsivity. Correct responses were rewarded with 4% stevia-water solution and a concomitant reinforcement sound, while incorrect responses were followed by an unpleasant sound (400 ms, 300 Hz square-wave) and a 1-3 second timeout period. A middle-grey background was displayed during non-stimulus periods. In a given trial, every stimulus had an equal probability of being presented, but with the constraint that a maximum of three consecutive presentations of the same visual class was allowed to prevent the development of a bias towards one of the response ports. Each training/test sessions lasted 40-60 minutes (*21, 22*). Stimulus presentation, response collection, and reward delivery were controlled by the MWorks software (`https://mworks.github.io`).

### Experimental design

**Training.** The animals were trained according to the following phases.

*Habituation*: the rats were first habituated to the operant chamber and were gradually trained to initiate trials autonomously and receive rewards from the lateral feeding needles. Initially, the animals were manually guided to the nose-poke ports, often reinforced with honey drops. A trial started once the animal touched the central port with its nose, signaled by a positive reinforcement tone, and followed by stevia-water rewards from both lateral feeding needles. This phase lasted about 7 days, progressing to the next phase once the animals initiated at least 100 trials autonomously in three consecutive sessions.

*Phase 1*: rats were conditioned to associate rewards with the presentation of paired auditory and visual stimuli. Initially, to facilitate the task, the visual stimuli were those with the lowest (0.25 Hz) and highest (4 Hz) temporal frequencies and were presented at full contrast alongside the fixed amplitude white noise sound for 2 seconds. Concomitantly, the reward was delivered from the correct response port, based on the class the visual stimulus belonged to. Advancement to the next phase was based on qualitative inspection of the behavior, ensuring that rats attended to the visual stimuli. Importantly, here, as in the rest of the training and test phases, only the visual stimuli were associated to a given reward port (i.e., to a given response class). The auditory stimuli were always task-irrelevant, i.e., were uninformative about the correct classification of the visual stimuli.

*Phase 2*: in this phase, the rats were actively reinforced based on their classification of the visual stimuli, presented again with a temporal frequency of either 0.25 or 4 Hz. That is, in this phase, the reward was delivered only after the rat had approached and licked the correct response port, while, in case of incorrect choice, the disturbing sound was delivered and the time-out period started. Again, both visual stimuli were paired with the same, uninformative fixed amplitude sound. Advancement to the next phase required that an animal correctly classified at least 70% of the visual stimuli in at least two consecutive sessions. Two of the 12 rats failed to meet this criterion and were excluded from further testing.

*Phases 3, 4, and 5.*: here, we gradually introduced the additional temporal frequencies of the visual stimuli. First, we added the gratings at 0.82 and 3.41 Hz (*Phase 3*), then those at 1.32 and 2.91 Hz (*Phase 4*), bringing to six the total of stimuli to classify. Finally, we presented all nine possible visual temporal frequencies (*Phase 5*). Also in this case, each advancement to the next phase depended on whether an animal was able to achieve an accuracy criterion of 70% correct responses in at least two consecutive training sessions. The 2.12 Hz stimulus, serving as an arbitrary boundary between the "low temporal frequency" and "high temporal frequency" classes, was randomly assigned to one of the classes in each trial and rewarded accordingly.

Upon achieving 70% accuracy in at least two consecutive sessions, indicating proficiency in classifying all visual stimuli, we progressively reduced the contrast of the visual stimuli. Previous studies have shown that A1 impacts V1 more strongly when auditory stimulus power is high (>70 dB SPL) and visual stimulus contrast is low (*13, 17*). The contrast reduction was performed in four steps based on accuracy criteria. If a rat maintained high accuracy (70% correct choices) for at least two consecutive training days at a given contrast level, the contrast was further decreased. The sequence of contrast reduction was 75%, 50%, 35%, and finally 25% of full contrast. The learning trajectory during the training phase is illustrated in Fig. S3 for two representative rats and typically lasted between 55 and 100 daily sessions.

**Test phase.** After completing the training phase, the rats entered the test phase. Here, the visual stimuli presented in each trial could be paired not only with the fixed amplitude sound (as done during training) but also with the amplitude modulated auditory stimuli. Specifically, the trials in the test phase fell into four main categories, based on the combination of visual and auditory stimuli (see Fig. 1B).

- In the "V + fixed amplitude sound" trials, which matched those in training Phase 5, the visual gratings of varying temporal frequencies were paired with the fixed amplitude white noise sound (green cells in Fig. 1B).
- In the "V + AM sounds" trials, the visual gratings were paired with each of the amplitude modulated sounds at the various frequencies, resulting in all 81 possible pairwise combinations of visual and auditory frequencies (cells in the squared matrix of Fig. 1B).

- In the “V only” or unimodal visual trials, the drifting gratings were presented without any accompanying sound.
- In the “A only” or unimodal auditory trials, the amplitude modulated sounds were presented alone, without the visual gratings. Since the rats had never been trained to classify the auditory stimuli, we simply collected their spontaneous responses to these sounds, without providing any feedback (i.e., neither reward nor time-out) about their choices.

Again, throughout both the training and test phases, the temporal frequency of the visual stimuli remained the only task-relevant feature for inferring the correct reward location. In contrast, the auditory stimuli were always task-irrelevant and did not indicate the reward side. We verified that the rats did not implicitly learn to classify the auditory stimuli according to their temporal frequency, by plotting their choices in the “A only” trials (Fig. S4). The resulting group average psychometric curve was flat, showing no modulation as a function of sound frequency.

**Sound stimuli generation and estimation of sound intensity**

We measured the intensity of the sound stimuli with a phonometer (Tadeto SL720). At any point in time, the phonometer provides an instantaneous estimate of the sound intensity based on a characteristic sampling window (0.125 s). To avoid any interactions between the temporal modulation of our sound stimuli and the finite sampling rate of the phonometer, we devised and calibrated a transformation that estimated the instantaneous sound intensity as a function of the nominal control signal. This transformation takes into account that the control signal $a(t)$ is converted into a sound pressure wave $p(t)$ by the amplifier/speaker, assuming minimal distortion (i.e., that there is a linear relationship between $a$ and $p$), and that the measurements of sound intensities $L(t)$ returned by the phonometer are in dB - i.e., they are a nonlinear function of $p(t)$. We note here that our dB measurement are done using “A” frequency weighting, as in previous studies (*14*).

To carry out this computation, we first verified that the relationship between the nominal control signal $a(t)$ and $p(t)$ as measured by the phonometer was linear. To do so, we measured with the phonometer the intensity $L$ of a uniform white noise sequence lasting 10 seconds. The long duration ensured that we got a precise estimate of the intensity in dB. We repeated the measurements 10 times, each time dimming the control signal $a$ by a factor of 0.8. All these measures were taken by placing the phonometer inside the sound-proof operant chambers, in the same position where the head of the rats was located during the task. Under the same settings, we also measured the intensity $L_a$ of the ambient noise inside the operant chambers (32 dB). The relationship between the sound intensity $L$ (in decibels) of a white noise sequence and its sound pressure level $p$ (in Pascals) is given by:

$$L = 10 \times \log_{10}\left(\frac{\bar{p}^2 + p_a^2}{p_0^2}\right) \tag{S1}$$

where $\bar{p}$ is the root-mean-square (RMS) of $p(t)$ over a time window $\tau$, $p_0 = 20\ \mu$P is the reference pressure, and $p_a$ is the pressure of the ambient noise. Inverting the equation, we can compute the sound pressure from the intensity measurements. The ambient sound pressure is given by:

$$p_a = p_0\sqrt{10^{\frac{L_a}{10}}} \tag{S2}$$

The sound pressure of the white noise sequences is, therefore:

$$p = p_0\sqrt{10^{\frac{L}{10}} - \left(\frac{p_a}{p_0}\right)^2} \tag{S3}$$

We could thus compute $p$ from the measured values of $L$, finding that, as expected under the assumption of a linear mapping from $a$ to $p$, the average pressure for each sequence was $0.8 \pm 0.01$ times as intense as the previous one. This confirmed that $p(t) = f \times a(t)$ for some value of $f$ and allowed us to estimate the factor $f$ of the linear relationship as $f = 0.39$ P. We repeated the same procedure with gaussian white noise sequences, confirming the result and obtaining a compatible estimate for $f$. We could then use the estimated value of $f$ to calculate the sound intensity (in dB) of all sound stimuli used in the experiment. For the fixed amplitude, white noise sound $a_{fix}(t)$, this results in $L_{fix} = 74.8$ dB. For the amplitude modulated stimuli, their sound intensity profile $L_{AM}(t)$ is in principle influenced by the duration $\tau$ of the time window over which the RMS of $p_{AM}(t)$ is computed. For short values of $\tau$, close to the white-noise-sampling period ($T_{WN} = \frac{1}{\nu_{WN}}$), noise fluctuations dominate the temporal profile. By contrast, long $\tau$s, comparable to the period of the modulation, will effectively smooth the modulation, resulting in $L_{AM}$ values that do not accurately represent the dynamic characteristics of the signal. For a wide range of intermediate time windows, however, the computed sound intensity is independent of the details of the fluctuation of the carrier and its temporal profile correctly tracks the temporal modulation.

More specifically, the RMS of $p_{\mathrm{AM}}(t)$ over a time window $\tau$ is defined as

$$\bar{p}_{\mathrm{AM}}(t)_\tau = \sqrt{\left\langle f^2 \cdot a_{\mathrm{AM}}(t)^2 \cdot m(t)^2 \right\rangle_\tau} \tag{S4}$$

If $\tau$ is smaller than the temporal scale of the modulation, then $m(t)$ is approximately constant inside the integration window and

$$\left\langle f^2 \cdot a_{\mathrm{AM}}(t)^2 \cdot m(t)^2 \right\rangle_\tau \simeq f^2 m(t)^2 \left\langle a_{\mathrm{AM}}(t)^2 \right\rangle_\tau \tag{S5}$$

On the other hand, if $\tau$ is larger than the sampling rate of the white noise ($\tau \gg T_{\mathrm{WN}}$) there will be enough samples in the window for the RMS of $a_{\mathrm{AM}}(t)$ to be equal to the uniform distribution average:

$$\bar{a}^2_{\mathrm{AM}} = \left\langle a_{\mathrm{AM}}(t)^2 \right\rangle_\tau \simeq \mathbb{E}[a_{\mathrm{AM}}(t)^2]_{p(a_{\mathrm{AM}})} = \int_{-1}^{1} \frac{1}{2} a^2 \, da = \frac{1}{3} \tag{S6}$$

Therefore, the resulting temporal intensity profile of the AM sounds (in dB), when using one of the intermediate values of $\tau$ can be computed as:

$$L_{\mathrm{AM}}(t) = 10 \log_{10} \left( \left( \frac{f \cdot \bar{a}_{\mathrm{AM}} \left( \frac{1 - cos(\omega t)}{2} \right)}{p_0} \right)^2 + \left( \frac{p_{\mathrm{a}}}{p_0} \right)^2 \right) \tag{S7}$$

The sound intensity profiles $L_{\mathrm{AM}}(t)$ for each modulation frequency (computed with $\tau = 5$ ms $\simeq 220 T_{\mathrm{WN}}$), as well as the profiles of $L_{\mathrm{fix}}$ and $L_{\mathrm{a}}$ (with the latter corresponding to the "visual only" condition) are shown in Fig. S2B.

To estimate the sound intensity experienced by a rat during each trial, we computed the average intensity $\bar{L}_{\mathrm{n}}(T)$ of the trial condition $n$ throughout a specific time window with duration $T$. This duration was equal to the *reaction time* ($RcT$) of the animal in that given trial, i.e., to the period during which the rat was exposed to the sound stimulus before initiating a motor response. The reaction time is defined as:

$$RcT = RT - MoRsT \tag{S8}$$

where $RT$ is the *response time* (i.e., the time from the trial onset to the moment a response port is reached), and $MoRsT$ is the *motor response time* (i.e., the time taken by the animal to reach the selected response port, after leaving the central port for the first time). In the recorded data, only the response time was directly measured for each trial. To estimate the reaction time, we assigned to $MoRsT$ a fixed value across all trials, as determined in one of our previous studies, where we carefully mapped the trajectories and durations of rat head movements in a similar experimental setup (*24*). Specifically, we used $MoRsT = 0.3$ s, which corresponds to the duration of a direct ballistic movement from the central port to either response port. This type of movement was the most frequently observed in that study (1149 out of 1359 recordings). Hence, for each trial $i$, we computed the average perceived intensity as $\bar{L}_{\mathrm{n}}(RcT_i) = \bar{L}_{\mathrm{n}}(RT_i - MoRsT)$. An example of the computation is shown on the "0.25 Hz AM sound" row of Fig. S2B.

The resulting values for each trial are shown as scatter plots in Fig. S2C, along with violin plots to illustrate their distribution in each sound condition for every individual rat (the dashed lines report the rat group average intensities). The averages of these sound intensities obtained for each rat are those reported on the x axis of Fig. 2C (squares, triangles, and crosses). Note, however, that, in that figure, the AM conditions with modulation frequencies $1.32\ \mathrm{Hz} \leq \mathrm{TF} \leq 4\ \mathrm{Hz}$ have been grouped together, given that their average intensity was virtually identical (as shown in Fig. S2C).

#### 2-way ANOVA with repeated measures analysis

To perform the two-way ANOVA with repeated measures, we first calculated the probabilities of each rat to classify a visual stimulus as belonging to the "High TF" category across all experimental conditions. This resulted in 10 data points for each of the stimulus conditions analyzed in Fig. 1C and D, corresponding to the 10 rats in our study. The ANOVA analysis requires two key assumptions for the data distributions:

1. Normality, i.e., the data should follow a Gaussian distribution. This is particularly important for probability distributions, as distributions near 0 and 1 can saturate and deviate from normality. To address this, we transformed the psychometric data using the arcsine of the square root of each value. After the transformation, we tested the normality of the distributions with the Shapiro-Wilk test (*25*) and we verified that it was satisfied by each data point distribution.

2. Homoscedasticity, i.e., the variance should be equal across experimental conditions. The arcsine transform also helps stabilize the variance for proportional data. This was assessed using Levene's test. Following the transform, the test could not reject the null hypothesis

that all distributions had equal variance ($p = 0.3421$), indicating that the assumption of homoscedasticity was satisfied.

We then proceeded to calculate the two-way ANOVA with repeated measures. Our experimental design involved two independent variables: the TF of the visual stimuli, with 9 levels, and the experimental condition, with 3 levels. The latter were as follows. For the analysis carried out in Fig. 1C: the "V + fixed amplitude sound", the "V + AM sounds (congruent)" and the "V + AM sounds (anti-congruent)"; for the analysis carried out in Fig. 1D: the "V + fixed amplitude sound", the "V + AM sounds" and the "V only". The dependent variable was rat response, measured as the probability of classifying a visual stimulus in the "High TF" category for each experimental condition, which was recorded for the same 10 animals across all the different levels of the independent variables. Table 1a and 1b (top half) present the results of this analysis for the data shown, respectively, in Fig. 1C and D, reporting, in both cases, significant effects for both the visual TF and the experimental condition variables, as well as a significant interaction between them. This indicates that the dependent variable changed across different visual TFs and experimental conditions, and that there was a combined influence of visual TF and experimental condition on the dependent variable.

Finally, to determine which specific levels of the experimental conditions contributed to the main effect reported in the ANOVA table, we conducted post-hoc tests. We performed pairwise comparisons of the different levels of the experimental conditions and evaluated their effects using the Tukey test, as summarized in the bottom halves of Table 2a and 2b.

### Ideal Observer Model

In this section we derive the ideal observer model used to predict the responses of the rats in the visual rate classification task. The ideal observer follows a standard structure for a sensory classification task [see e.g. (*26*)], with the added hypothesis that the measurement $x$ (an internal, abstract representation of the visual stimulus) is scaled by a gain factor $\gamma$ which can depend on the intensity of the auditory stimulus. This hypothesis stands on its own on an abstract level, but we show below that it can be derived as the consequence of a putative suppressive effect of sound on the activity of visual cortical neurons.

On each trial of the two-alternative forced choice (2AFC) task, the visual stimuli were assigned a temporal frequency which was randomly chosen from a pool of 9 possible values (0.25, 0.82, 1.32, 1.75, 2.12, 2.48, 2.91, 3.41 and 4 Hz). In the following, we will denote this visual temporal frequency by $s$, for stimulus. The goal of the animal was to accurately classify each visual stimulus as belonging to the low frequency class ($L$, for gratings with frequencies from 0.25 to 2.12 Hz) or to the high frequency class ($H$, from 2.12 to 4 Hz). Each of the 9 frequency values had an equal chance of being presented in a trial. The central value was randomly assigned to the $H$ or $L$ class, and therefore also $H$ and $L$ had an equal chance of being presented. The probabilistic structure of the task can also be restated as follows. On each trial, the stimulus class $C$ is chosen with equal chance to be $H$ or $L$, that is

$$p(C = H) = p(C = L) = \frac{1}{2} \tag{S9}$$

Based on the class, the visual stimulus $s$ presented to the animal is selected with probability

$$p(s|C) = \frac{\delta(s - s_0)}{2K + 1} + \frac{2}{2K + 1} \sum_{k=1}^{K} \delta(s - s_k^C) \tag{S10}$$

where

$$s_0 = 2.12 \tag{S11}$$

$$s_k^L \in \{0.25; 0.82; 1.32; 1.75\} \tag{S12}$$

$$s_k^H \in \{2.48; 2.91; 3.41; 4\} \tag{S13}$$

$$K = 4 \tag{S14}$$

and for convenience with our derivations below we order the index $k$ symmetrically around $s_0$, such that $s_1^H = 2.48$, $s_2^H = 2.91$, etc and $s_1^L = 1.75$, $s_2^L = 1.32$, etc. Our task here is to compute the probability $P(\hat{C} = H \mid s, n)$ that the observer chooses the class "high visual frequency" ($\hat{C} = H$), when the frequency of the visual stimulus is $s$ and the sound condition is $n$.

We start by modeling the process that on each trial gives rise to the measurement $x$, given the visual stimulus $s$ and the sound condition $n$. We initially disregard the effect of the sound, which we will add back later. We assume that a stimulus $s$ gives rise to a certain pattern $\vec{r}$ of firing rates in V1, where $r_i$ is the rate of the $i$-th neuron. The population response is normally distributed around a stimulus-dependent value $\vec{\mu}(s)$: $\vec{r} \sim \mathcal{N}(\vec{\mu}(s), \Sigma)$, where $\Sigma$ is a generic (but stimulus-independent) covariance matrix.

The subject generates an internal representation (or *measurement*) $x$ of the external stimulus $s$ based on the activation pattern $\vec{r}$. We assume that this measurement is a projection of the vector of activities $\vec{r}$ on some measurement axis $y = \vec{u} + t\vec{v}$; in other words,

$$x = (\vec{r} - \vec{u}) \cdot \vec{v} \tag{S15}$$

This is biologically plausible if, for example, the measurement $x$ is the activity of a readout neuron with connectivity strength $v_i$ with each neuron coding $s$. Ideally, the measurement axis $y$ used by the subject is the one that allows to best discriminate the stimulus classes, therefore allowing for the best performance, but for the purpose of developing the model we do not need to make any particular assumptions on $\vec{u}$ and $\vec{v}$. Because $x$ is an affine transformation of a normally-distributed random variable, it is itself a normally distributed random variable:

$$x \sim \mathcal{N}\left((\vec{\mu}(s) - \vec{u}) \cdot \vec{v}, \vec{v}^{\intercal}\Sigma\vec{v}\right) \tag{S16}$$

If the projection operation is itself noisy and introduces additional Gaussian noise of standard deviation $\tau$, the distribution of the measurement becomes simply

$$x \sim \mathcal{N}\left((\vec{\mu}(s) - \vec{u}) \cdot \vec{v}, \vec{v}^{\intercal}\Sigma\vec{v} + \tau^2\right) \tag{S17}$$

In the following, we will lump both sources of variability into a single parameter $\rho$,

$$\rho^2 = \vec{v}^{\intercal}\Sigma\vec{v} + \tau^2 \tag{S18}$$

We will conceptualize $\rho$ as the general level of sensory noise present in the system for the purposes of this task. The distribution of the measurement will therefore be

$$x \sim \mathcal{N}\left((\vec{\mu}(s) - \vec{u}) \cdot \vec{v}, \rho^2\right) = \mathcal{N}\left(\nu(s), \rho^2\right) \tag{S19}$$

where we have defined

$$\nu(s) = (\vec{\mu}(s) - \vec{u}) \cdot \vec{v} \tag{S20}$$

for notational convenience.

To decide if a stimulus has high frequency ($H$) or low frequency ($L$), we assume that the subject performs Bayesian inference to compute a posterior probability over the classes $H$ and $L$ given the measurement and reports the class with highest posterior. In our case (see below for a detailed justification), this will be equivalent to finding a decision boundary $x^*$ such that $P(H \mid x^*) = P(L \mid x^*)$, and reporting $H$ if $x > x^*$ and $L$ if $x < x^*$. On average over the distribution of the measurement given the stimulus, the probability of reporting a given class given the stimulus (which we may also call the psychometric function) will then be

$$p(\hat{C} = H \mid s) = \int_{x:p(H|x)>p(L|x)} p(x \mid s)dx \tag{S21}$$

$$= p(x > x^* \mid s) \tag{S22}$$

$$= \int_{x^*}^{\infty} p(x \mid s)dx \tag{S23}$$

and $p(\hat{C} = L \mid s) = 1 - P(\hat{C} = H \mid s)$. We will now calculate $x^*$ and consequently we will derive $p(\hat{C} = H|s)$.

Let's consider the simplest possible scenario for $\vec{\mu}$, namely the case where $\vec{\mu}$ is linear in $s$:

$$\vec{\mu}(s) = \vec{\alpha} + \vec{\beta}s \tag{S24}$$

In this case, the mapping between $s$ and $x$ also becomes linear:

$$\nu(s) = (\vec{\mu}(s) - \vec{u}) \cdot \vec{v} = \vec{\alpha} \cdot \vec{v} + (\vec{\beta} \cdot \vec{v})s - \vec{u} \cdot \vec{v} = a + bs \tag{S25}$$

with $a = (\vec{\alpha} - \vec{u}) \cdot \vec{v}$ and $b = \vec{\beta} \cdot \vec{v}$.

We will now show that when $\vec{\mu}$ is linear the symmetry of the problem and the shape of the response distributions imply that $x^*$ is unique and $x^* = \nu(s_0)$. If we compute the posteriors explicitly from the class-conditional stimulus distributions in Equation S10 we get

$$p(H|x) - p(L|x) = \frac{p(x|H)p(H)}{p(x)} - \frac{p(x|L)p(L)}{p(x)} \tag{S26}$$

$$\propto p(x \mid H) - p(x \mid L) \tag{S27}$$

$$= \frac{p(x|s_0)}{2K+1} + \frac{2}{2K+1}\sum_{k=1}^{K} p(x|s_k^H) - \frac{p(x|s_0)}{2K+1} - \frac{2}{2K+1}\sum_{k=1}^{K} p(x|s_k^L) \tag{S28}$$

$$\propto \sum_{k=1}^{K} p(x|s_k^H) - p(x|s_k^L) \tag{S29}$$

$$\propto \sum_{k=1}^{K} \exp\left[-\frac{(x - \nu(s_k^H))^2}{2\rho^2}\right] - \exp\left[-\frac{(x - \nu(s_k^L))^2}{2\rho^2}\right] \tag{S30}$$

$$= \sum_{k=1}^{K} \exp\left[-\frac{(x - a - bs_k^H)^2}{2\rho^2}\right] - \exp\left[-\frac{(x - a - bs_k^L)^2}{2\rho^2}\right] \tag{S31}$$

Note now that the symmetry of the class-conditional stimulus distributions in Equation S10 is such that

$$\begin{cases} (s_0 - s_k^H)^2 = (s_0 - s_k^L)^2 \; \forall k \\ (s - s_k^H)^2 < (s - s_k^L)^2 \; \forall k \quad \text{if } s > s_0 \\ (s - s_k^H)^2 > (s - s_k^L)^2 \; \forall k \quad \text{if } s < s_0 \end{cases} \tag{S32}$$

Also, for any value of $x$ and $s$,

$$x - \nu(s) = a - b\nu^{-1}(x) - a - bs = b(\nu^{-1}(x) - s) \tag{S33}$$

which means that, if $b > 0$ (and simply reversing the inequalities if $b < 0$),

$$\begin{cases} (x - \nu(s_k^H))^2 = (x - \nu(s_k^L))^2 \; \forall k \\ (x - \nu(s_k^H))^2 < (x - \nu(s_k^L))^2 \; \forall k \quad \text{if } x > \nu(s_0) \\ (x - \nu(s_k^H))^2 > (x - \nu(s_k^L))^2 \; \forall k \quad \text{if } x > \nu(s_0) \end{cases} \tag{S34}$$

Therefore,

$$\begin{cases} p(H|x) = p(L|x) \quad \text{if } x = \nu(s_0) = a + bs_0 \\ p(H|x) > p(L|x) \quad \text{if } x > \nu(s_0) = a + bs_0 \\ p(H|x) < p(L|x) \quad \text{if } x < \nu(s_0) = a + bs_0 \end{cases} \tag{S35}$$

Thus as a consequence of the symmetry of the problem the decision boundary $x^*$ is simply

$$x^* = \nu(s_0) = a + bs_0 = (\vec{\alpha} + \vec{\beta}s_0 - \vec{u}) \cdot \vec{v} \tag{S36}$$

Hence:

$$p(\hat{C} = H \mid s) = p(x > x^* \mid s) = \int_{x^*}^{\infty} p(x \mid s) dx \tag{S37}$$

$$= \int_{x^*}^{\infty} \frac{1}{\sqrt{2\pi\rho^2}} \exp\left[-\frac{(x - (\vec{\mu}(s) - \vec{u}) \cdot \vec{v})^2}{2\rho^2}\right] dx \tag{S38}$$

$$= \Phi\left[\frac{(\vec{\mu}(s) - \vec{u}) \cdot \vec{v} - x^*}{\rho}\right] \tag{S39}$$

$$= \Phi\left[\frac{(\vec{\mu}(s) - \vec{u}) \cdot \vec{v} - (\vec{\mu}(s_0) - \vec{u}) \cdot \vec{v}}{\rho}\right] \tag{S40}$$

$$= \Phi\left[\frac{(\vec{\mu}(s) - \vec{\mu}(s_0)) \cdot \vec{v}}{\rho}\right] \tag{S41}$$

$$= \Phi\left[\frac{\vec{\beta}(s - s_0) \cdot \vec{v}}{\rho}\right] \tag{S42}$$

$$= \Phi\left[\frac{s - s_0}{\sigma}\right] \tag{S43}$$

with

$$\sigma = \frac{\rho}{\vec{\beta} \cdot \vec{v}} \tag{S44}$$

As shown in Fig. 2A, we model the effect of an external sound stimulus $n$ as a suppression of the activity of V1 neurons.This suppression may result from direct inhibition of V1 neurons by auditory neurons, but it could arise from indirect effects as well. We model it by multiplying the mean response $\vec{\mu}(s)$ of the neurons by a sound-dependent factor $\gamma_n$, while keeping fixed the decision boundary $x^*$ learned during the training sessions. By substituting $\vec{\mu}(s)$ with $\gamma_n\vec{\mu}(s)$ in the integral above, we get:

$$p(\hat{C} = H \mid s, n) = \Phi\left[\frac{(\gamma_n\vec{\alpha} + \gamma_n\vec{\beta}s - \vec{u}) \cdot \vec{v} - x^*}{\rho}\right] = \Phi\left[\frac{\gamma_n s - s_0 + (\gamma_n - 1)\lambda}{\sigma}\right] \tag{S45}$$

with

$$\lambda = \frac{\vec{\alpha} \cdot \vec{v}}{\vec{\beta} \cdot \vec{v}} \tag{S46}$$

Now, if the circuit computing $x$ is adapted to extract the maximum amount of information from $\vec{r}$ in this task, at least in the case where the noise for $\vec{r}$ is isotropic, $\vec{v}$ will be parallel to $\vec{\beta}$ (this may not be true in presence of nontrivial noise correlation structure). On the other hand, if $\vec{\alpha}$ is a random high dimensional vector with no particular relation to $\vec{v}$, we can assume that with high probability

$\vec{\alpha} \cdot \vec{v} = 0$. Therefore, under these simple assumptions, $\lambda = \vec{\alpha} \cdot \vec{v} / \vec{\beta} \cdot \vec{v} = 0$. In this case, the expression for the probability of reporting $H$ in presence of auditory stimulus $n$ reduces to

$$p\left(\hat{C} = H \mid s, n\right) = \Phi\left[\frac{\gamma_n s - s_0}{\sigma}\right] \tag{S47}$$

where $\gamma_n$ and $\sigma$ are free parameters of the model. Finally, we note that while the linearity assumption for $\vec{\mu}(s)$ made in this section can seem restrictive, the derivations above remain valid as long as $\vec{\mu}(s) \cdot \vec{v}$ is (approximately) linear over the range of values of $s$ that are relevant in the task.

**Lapse rates.** We extended the ideal observer to account for lapses by defining two lapse rates $\epsilon_H$ and $\epsilon_L$, such that on each trial the observer has a probability $1 - \epsilon_H - \epsilon_L$ of actually engaging in the task, a probability $\epsilon_H$ of reporting $H$ independently of the stimulus, and a probability $\epsilon_L$ of reporting $L$ independently of the stimulus. Therefore,

$$p(\hat{C} = H \mid s, n) = \epsilon_H + (1 - \epsilon_H - \epsilon_L)\, \Phi\left[\frac{\gamma_n s - s_0}{\sigma}\right] \tag{S48}$$

and

$$p(\hat{C} = L \mid s, n) = 1 - p(\hat{C} = H \mid s, n) \tag{S49}$$

**Inferring the model parameters from experimental data**

We used the ideal observer model derived above to describe the data from our experiment. According to the formula for $p(\hat{C} = H \mid s, n)$, the behavior of each rat $r$ is characterized by some parameters $\theta^r = \left\{\sigma^r, \left\{\gamma_n^r\right\}, \epsilon_L^r, \epsilon_H^r\right\}$. We inferred the value of these parameters using a hierarchical Bayesian approach (*27,28*), for which we now give a brief conceptual overview (refer to Fig. S5 for a graphical depiction).

Denote by $D$ the set of recorded behavioral data, with $D_t^r$ being the choice ($H$ or $L$) of rat number $r \in \{1, \ldots, R\}$ on trial number $t \in \{1, \ldots, T_r\}$. We start by defining, for any rat, a prior probability for the parameters:

$$p(\theta^r) = p(\sigma^r) p(\epsilon_L^r, \epsilon_H^r) \prod_n p(\gamma_n^r) \tag{S50}$$

We also define a likelihood function for the parameters $\theta^r$. The likelihood is simply the probability of observing the rat choices $D^r = \{D_t^r\}_{t=1}^{T_r}$, given a certain value of $\theta^r$ and of the visual and auditory stimuli provided to the rat on all trials (respectively, $s^r = \{s_t^r\}_{t=1}^{T_r}$ and $n^r = \{n_t^r\}_{t=1}^{T_r}$), seen as a function of $\theta^r$:

$$\mathcal{L}^r(\theta^r) = p(D^r|\theta^r; s^r, n^r) \tag{S51}$$

$$= \prod_{t=1}^{T_r} p\left(\hat{C} = D_t^r \mid \theta^r; s_t^r, n_t^r\right) \tag{S52}$$

$$= \prod_{t:D_t^r=H} p\left(\hat{C} = H \mid \theta^r; s_t^r, n_t^r\right) \prod_{t:D_t^r=L} p\left(\hat{C} = L \mid \theta^r; s_t^r, n_t^r\right) \tag{S53}$$

where $p(\hat{C} = H|s, n)$ and $p(\hat{C} = L|s, n)$ are given by Equation S48 and Equation S49. The likelihood function therefore encapsulates the ideal observer model, developed in the previous section. By applying Bayes' theorem, we compute a joint posterior probability distribution for the parameters:

$$p(\theta^r \mid D^r) \propto p(D^r \mid \theta^r)p(\theta^r) \tag{S54}$$

(from now on, for convenience, we will omit the explicit dependency on $s$ and $n$ in the likelihood). However, since the rats belong to the same population and are subjected to the same experimental conditions, we assume that the parameters describing different animals will be related. In other words, gathering information about one rat tells us something about that specific rat, but should also inform us on what to expect about other rats, or rat behavior in general. We can build this assumption into the model by imposing that the parameters associated to different rats are all sampled from the same distribution, leaving the parameters of this higher-level probability distribution to be also inferred from the data. Besides extracting "population-level" information from the data of each rat, this procedure will also help constraining the rat-level parameters to reasonable values given what we observed in the other rats, thus acting as a regularizer for the inference process. More formally, our posterior will be:

$$p(\theta, \eta \mid D) \propto p(D \mid \theta)p(\theta \mid \eta)p(\eta) \tag{S55}$$

where $\theta = \{\theta^r\}_{r=1}^R$ are the rat specific parameters for all $R$ rats, and $\eta$ are the common (population) parameters.

More concretely, the priors we assign to the parameters ($p(\theta \mid \eta)$ in Equation S55) are:

$$(\epsilon_L^r, \epsilon_H^r, 1 - \epsilon_L^r - \epsilon_H^r) \sim \text{Dirichlet}(1, 1, 1) \quad \forall r \tag{S56}$$

$$\sigma^r \sim \text{Gamma}(k_\sigma, \theta_\sigma) \quad \forall r \tag{S57}$$

$$\gamma_n^r \sim \mathcal{N}(\mu_{\gamma_n}, \sigma_{\gamma_n}^2) \ \forall r, n \tag{S58}$$

It follows that our population parameters $\eta$ are the mean $\mu_{\gamma_n}$ and the standard deviation $\sigma_{\gamma_n}$ of the $\gamma_n$ parameters, and $k_\sigma$ and $\theta_\sigma$ for the perceptual noise. Following standard practice (*27, 29*), we assign weakly informative priors $p(\eta)$ to the population-level parameters:

$$\mu_{\gamma_n} \sim \mathcal{N}(\mu = 0, \sigma^2 = 9) \quad \forall n \tag{S59}$$

$$\sigma_{\gamma_n} \sim \text{Exponential}(\text{scale} = 3) \quad \forall n \tag{S60}$$

$$k_\sigma \sim \text{Exponential}(3) \tag{S61}$$

$$\theta_\sigma \sim \text{Exponential}(3) \tag{S62}$$

Finally, in this hierarchical scheme we also need a global likelihood function ($p(D \mid \theta)$ in Equation S55) which we obtain simply as the product of the rat-level likelihoods:

$$\mathcal{L}(\theta) = p\left(D \mid \{\vec{\gamma}_r, \sigma_r, \vec{\epsilon}_r\}_{r=1}^R\right) \tag{S63}$$

$$= \prod_{r=1}^{R} \prod_{t=1}^{T_r} p\left(\hat{C} = D_t^r \mid \theta^r; s_t^r, n_t^r\right) \tag{S64}$$

**Sampling the posterior distribution.** We performed inference with a Markov Chain Monte Carlo (MCMC) method, using the NUTS algorithm (*30*), as implemented in PyMC version 5.2.0 (*31*). This method samples parameters from the posterior distribution $p(\theta \mid D)$: with a high enough number of samples, the distribution of the samples matches closely the distribution of the posterior. We sampled 4 independent chains of 2000 draws each, 1000 of which we used for tuning the algorithm and subsequently discarded, and 1000 as actual samples of the target posterior distribution. The target acceptance probability parameter for NUTS was set at 0.99, and no divergences were detected during sampling.

### Model Comparison

To verify that the derived model describes well the experimental data, we compared it with other models that could also reproduce the psychometric curves of the rats. To perform the comparison, we used a Leave-One-Out Cross-Validation (*LOO*) estimate of the Expected Log pointwise Predictive Density (*ELPD*) (*32*). LOO estimates how well, on average, the model describes data it hasn't been trained on. To do so, it computes how well a model trained on all the trials (across all animals) except one predicts the outcome of that trial, and averages this value over all the trials. The ELPD therefore automatically balances goodness of fit with model complexity, penalizing overly complex models that tend to overfit the noise in the data. More formally, the LOO estimate of the ELPD for a model $\mathcal{M}$ for which we can sample the posterior distribution with MCMC is defined as:

$$\mathrm{ELPD_{LOO}}(\mathcal{M}) = \sum_{i=1}^{N} \log\left(\frac{1}{S}\sum_{j=1}^{S} p_{\mathcal{M}}(d_i \mid \theta_{(j;D_{-i})})\right) \tag{S65}$$

where $N = \sum_{r=1}^{R} T_r$ is the total number of trials in the data, $S$ is the number of posterior samples in the Markov chain for model $\mathcal{M}$, $p_{\mathcal{M}}$ is the likelihood function of the model, $d_i$ is the data from the $i$-th trial and $\theta_{(j;D_{-i})}$ is the $j$-th posterior sample of a Markov chain generated from all data except $d_i$ (note that each posterior distribution sample, including $\theta_{(j;D_{-i})}$, is a high-dimensional vector containing one value for each parameter in the model). In Equation S65, the argument of the logarithm represents the likelihood of the model $\mathcal{M}$ for the datapoint $d_i$, averaged over the posterior distribution over $\theta$ (where this posterior is obtained without looking at datapoint $d_i$). Indeed, since the training of $\mathcal{M}$ results in a posterior distribution rather than a single parameter vector, all quantities related to $\mathcal{M}$ must be averaged over this distribution. For MCMC training, this average is computed as a sum over all sampled parameters, given that the samples follow the posterior distribution: $\frac{1}{S}\sum_{j=1}^{S} p_{\mathcal{M}}(d_i \mid \theta_{(j;D_{-i})}) \simeq \int p_{\mathcal{M}}(d_i \mid \theta)p(\theta \mid D_{-i})d\theta = \langle p_{\mathcal{M}}(d_i \mid \theta)\rangle_{p(\theta|D_{-i})}$. In practice, computing Equation S65 requires training the model $N$ times, one per trial: given the large number of trials in our experiment, we approximated the leave-one-out procedure with pareto-smoothed importance sampling (PSIS-LOO) (*32*).

Computing the $\mathrm{ELPD_{LOO}}$ for different models gives us a relative measure of how well they describe the data. As shown in the inset of Fig. 2B, the model with the highest ELPD is the one with 5 distinct $\gamma$ parameters, one for each sound condition whose power is in a certain range (see main text for details).

We also compared this model with a model that incorporates the additional $\lambda$ parameter discussed in the ideal observer derivation above (Equation S46). This parameter is included with an analogous hierarchical structure to that of $\sigma$ in the main model discussed above, such that each rat has its own $\lambda^r$, with

$$\lambda^r \sim \mathcal{N}(\mu_\lambda, \sigma_\lambda^2) \quad \forall r \tag{S66}$$

$$\mu_\lambda \sim \mathcal{N}(\mu = 0, \sigma^2 = 9) \tag{S67}$$

$$\sigma_\lambda \sim \text{Exponential}(3) \tag{S68}$$

This comparison allowed us to test the hypothesis that, to the extent that the assumptions about stimulus encoding made in that section are valid, the vector of resting activities of neurons $\vec{\alpha}$ has no effect on the projection of measurement. The resulting $(5\gamma 1\lambda 1\sigma)$ model performance is indeed comparable to the $(5\gamma 1\sigma)$ model performance ($\Delta$ ELPD = 0.74 ± 0.9; posterior mean ± std. err.). Moreover, the inferred population value of $\lambda$ in the $(5\gamma 1\lambda 1\sigma)$ model is consistent with 0: the posterior estimate is $\mu_\lambda = -0.17 \pm 0.19$ (mean ± std. err.), and the probability of direction is $p(\mu_\lambda < 0|D) = 0.821$. The probability of direction is the proportion of the posterior distribution for a parameter that has the same sign as the median, and in simple regression scenarios it is related to a frequentist two-sided p-value ($p \simeq 2(1 - pd)$) (*33*).

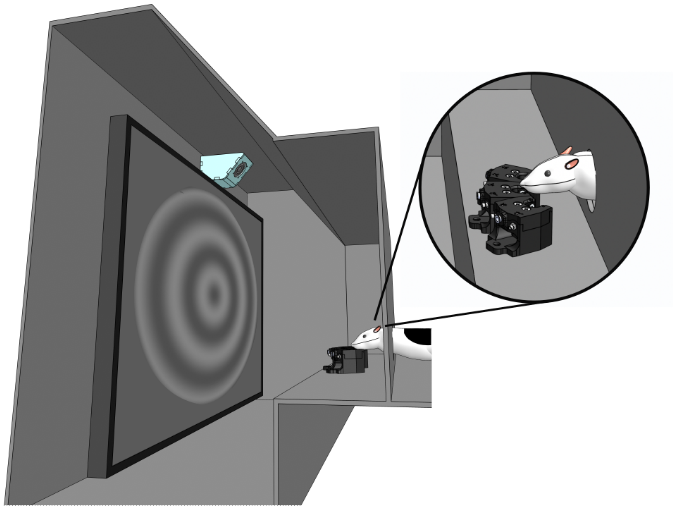

**Figure S1**: **Schematic of the operant chamber used to train the rats in the visual discrimination task.** The image depicts the rendering of a rat extending its head through a viewing hole in the wall of the chamber. The hole is placed in front of the stimulus display (featuring an example circular grating) and the speaker (in cyan) used to deliver the task-irrelevant sounds. The hole also allows access to an array of three nose-poke ports that the rat uses to trigger stimulus presentation (by licking the central port) and report the temporal frequency of the circular drifting gratings (left and right ports). The lateral ports are connected to computer-controlled syringe pumps (not shown in the image) for delivery of the liquid reward

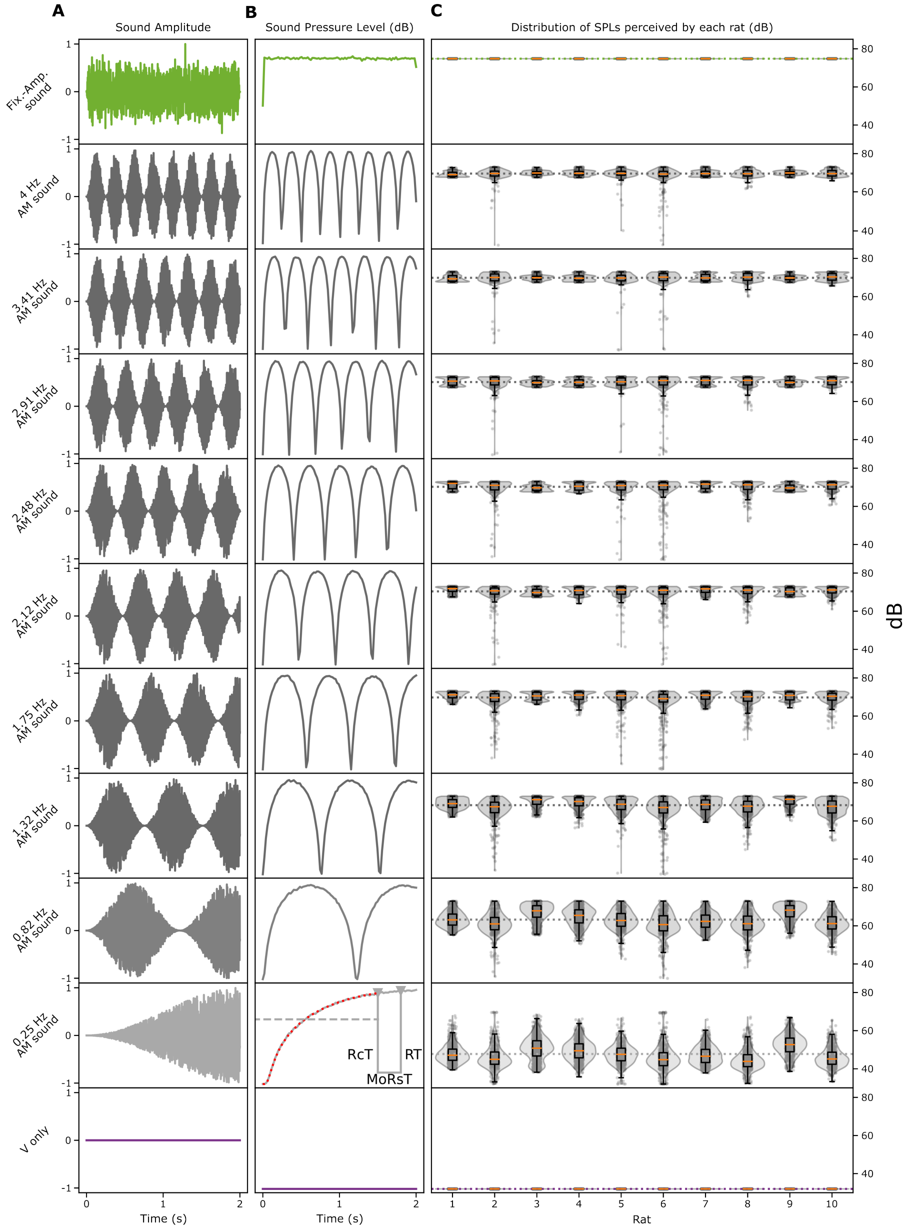


**Figure S2**: (Caption next page.)

**Figure S2**: **The auditory stimuli and their intensity.** Each row refers to a distinct auditory stimulus used in the experiment: 1) the white noise burst with fixed maximal amplitude (top row; green); 2) the amplitude modulated white noise bursts, with the 9 different temporal frequencies of the sinusoidal envelops (middle rows; gray); and 3) the absence of the auditory stimulus (bottom row; purple). **A**, The waveforms of the auditory stimuli used to drive the speakers that delivered the sounds to the rats (note that, for the sake of visualization, the sounds are not shown at their actual sampling frequency of 44.1 KHz but they have been downsampled to 500 Hz). **B**, The intensities of the auditory stimuli in dB as a function of time, as computed based on the waveforms shown in **A** and the measured intensity of 10 s long white noise bursts with different amplitude (*20*). The panel referring to the TF of 0.25 Hz illustrates how the average sound intensity experienced by a rat in a given trial was computed, based on the reaction time $RcT$ of the animal. The latter was obtained by subtracting the estimated motor response time $MoReT$ to the measured response time $RT$ (*20*). **C**, Distributions of the sound intensities experienced by each rat in every stimulus condition.

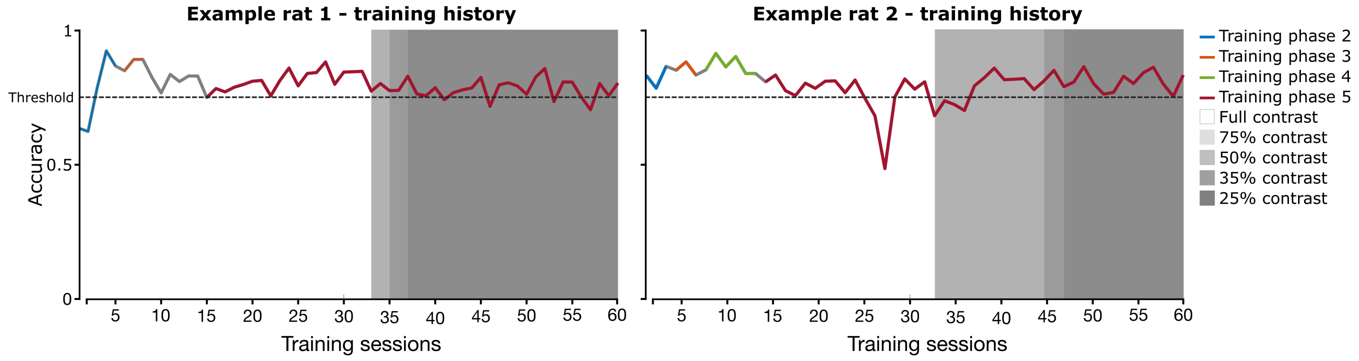


**Figure S3**: **Discrimination accuracy of two example rats across the different phases of the training procedure.** The lines show the percentage of correct choices of each rat in classifying the temporal frequency of the drifting gratings across 60 daily training sessions. The color of the line indicates the training phase (from 2 to 5; see caption), while the intensity of the gray patches denotes the various stages of contrast reduction performed in the final phase of the training (see caption). The dashed horizontal line represents the criterion accuracy (70%) required to progress to the next training phase. To advance, rats had to achieve performance above criterion for at least two consecutive days within the same training phase.

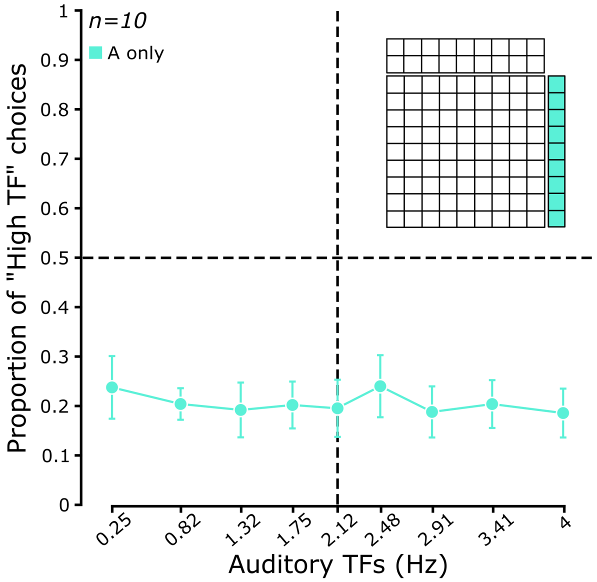


**Figure S4**: **Rats are not sensitive to the temporal frequency of the unimodal, purely auditory stimuli.** Group average proportion of "high TF" choices (n=10 rats) as a function of the TF of the unimodal auditory stimuli. Error bars are SEM.

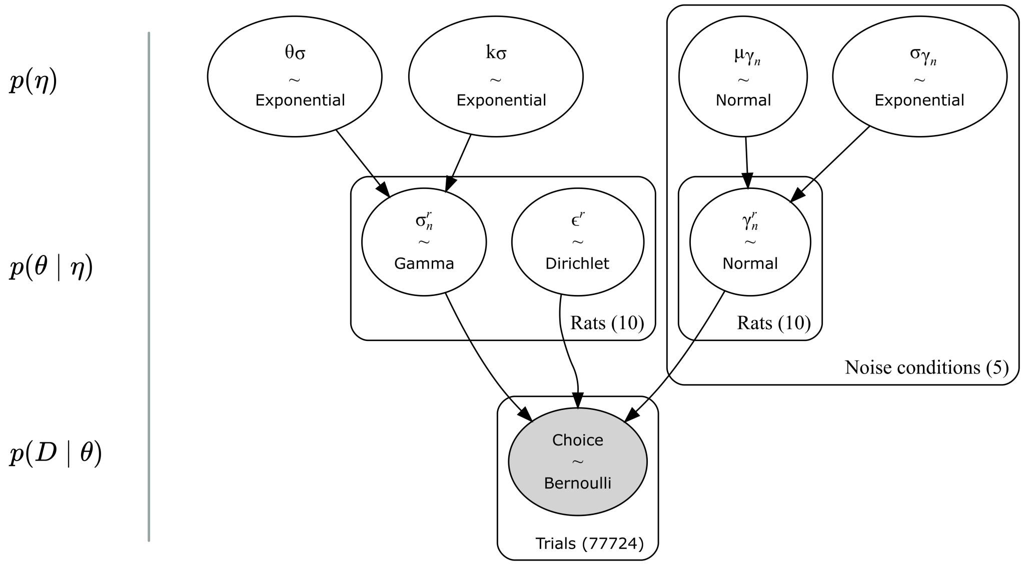


**Figure S5**: **Schematic of Bayesian hierarchical inference for our ideal observer model**. On the left, each row is labeled as representing the population parameters priors ($p(\eta)$), the subject-level parameters priors ($p(\theta \mid \eta)$) or the likelihood ($p(D \mid \theta)$). On the right, each bubble represents a parameter of the model. The name of the parameter and its distribution are reported. The rectangular plates indicate that multiple iid variables have been grouped. The group name and the corresponding number of variables are indicated on the bottom left of each plate. Each arrow represents the dependencies of the model. Gray coloring indicates that the variable is conditioned to observations.
